# mTOR repression in response to amino acid starvation promotes ECM degradation through MT1-MMP endocytosis arrest

**DOI:** 10.1101/2021.01.29.428784

**Authors:** Cecilia Colombero, David Remy, Sandra Antoine, Anne-Sophie Macé, Pedro Monteiro, Nadia ElKhatib, Margot Fournier, Ahmed Dahmani, Elodie Montaudon, Guillaume Montagnac, Elisabetta Marangoni, Philippe Chavrier

## Abstract

Under conditions of starvation, normal and tumor epithelial cells can rewire their metabolism towards the consumption of extracellular matrix-derived components as nutrient sources. The mechanism of pericellular matrix degradation by starved cells has been largely overlooked. Here we show that matrix degradation by breast and pancreatic tumor cells and patient-derived xenograft explants increases by one order of magnitude upon amino acid and growth factor deprivation. In addition, we found that collagenolysis requires the invadopodia components, TKS5 and the transmembrane metalloproteinase, MT1-MMP, which are key to the tumor invasion program. Increased collagenolysis is controlled by mTOR repression upon nutrient depletion or pharmacological inhibition by rapamycin. Our results reveal that starvation hampers clathrin-mediated endocytosis, resulting in MT1-MMP accumulation in arrested clathrin-coated pits. Our study uncovers a new mechanism whereby mTOR repression in starved cells leads to the repurposing of abundant plasma membrane clathrin-coated pits into robust ECM-degradative assemblies.

## Introduction

Metabolic reprogramming is a hallmark of cancer cells, which adapt their nutritional strategies to match their elevated metabolic needs ^1,2^. In certain microenvironments including in poorly perfused tumors, free nutrients such as amino acids (AAs) can be limiting, and extracellular proteins are used as alternative resources ^3,4^. In desmoplastic microenvironments in which tumors are encased such as in pancreatic and breast cancers, recent studies found that cancer cells can internalize proteolytic extracellular matrix (ECM) fragments, including peptides derived from fibronectin and type I collagen, which account for most of the extracellular biomass in these tumors ^5–7^. Degradation of ECM-derived peptides in lysosomes contributes to AA supply that fuels the tumor metabolism and supports tumor survival and proliferation ^6–8^. However, the mechanism underlying pericellular ECM breakdown under nutrient-depleted conditions is unknown.

The cell response to nutrients is controlled by the kinase mechanistic target of rapamycin (mTOR), which assembles into distinct protein complexes known as mTOR Complex 1 and 2 (mTORC1 and 2) ^9^. Only mTORC1 is sensitive to acute treatment by the anticancer drug, rapamycin ^9^. Under AA replete conditions, mTORC1 localizes to the lysosomes and phosphorylates several substrates including S6K and 4E-BP1, promoting protein translation and cell growth. Upon AA starvation, mTORC1 is inactivated inducing autophagy, cellular catabolism and translation shut-down ^9^.

The protease-dependent invasion program of tumor cells is mediated by invadopodia, which are F-actin-, cortactin-based cell-matrix contacts that enzymatically degrade and push confining ECM fibers aside to allow cell movement ^10–13^. The scaffolding protein, TKS5, plays a pivotal role in the assembly and surface accumulation of the trans-membrane matrix metalloproteinase and collagenase, MT1-MMP, to invadopodia ^13–16^.

Here, we investigated the mechanism of ECM degradation under conditions of nutrient scarcity in relation with mTOR signaling. Our findings uncover a novel mechanism that leads to the repurposing of the invadopodial MT1-MMP/TKS5 axis. This program, which is controlled by mTOR signaling, inhibits the endocytic clearance of MT1-MMP and triggers its accumulation in arrested plasma membrane clathrin-coated pits (CCPs) to actively degrade the collagen matrix in an AA-depleted environment.

## Results

### Starvation of tumor cells enhances invadopodia-mediated pericellular matrix degradation

MDA-MB-231 cells were selected as a model of breast cancer cells well known for producing robust invadopodia formation. When plated on fluorescently-labeled gelatin for 60 min in complete medium (CM), several gelatin degradation spots were visible underneath the cells, which coincided with TKS5-positive invadopodia (Figure 1A). The consequences of starvation on ECM degradation were assessed by culturing cells in AA- and serum-depleted medium (EBSS). The gelatinolytic activity of starved cells increased robustly overtime in relation with an extended punctate TKS5 pattern, and was strongly impaired by treatment with GM6001, a broad-spectrum MMP inhibitor (Figure 1A and E). Similarly, when cells were embedded within a 3D fibrillary collagen network, the main component of interstitial ECM tissue, and stained with Col1-¾C antibody that recorded collagen cleavage cumulated over the incubation period, we observed a strong enhancement of collagen cleavage by starved cells as compared to replete conditions (Figure 1B and F). Induction of collagen cleavage upon starvation was similarly observed in pancreatic adenocarcinoma Bx-PC3 cells (Supplemental Figure 1AB). MT1-MMP can be inhibited by tissue inhibitors of metalloproteinases (TIMPs). EBSS medium was supplemented with increasing amount of recombinant TIMP-2. Only at the dose of 2 μg/ml (*i.e.* 20-200-fold TIMP-2 concentration in tissues, biological fluids or in CM, not shown and ^17^), was recombinant human TIMP-2 capable of fully repressing collagenolysis (Supplemental Figure 1C). These data show that the absence of TIMP-2 in EBSS medium does not account for the observed changes in matrix degradation by starved cells.

**Figure 1.**
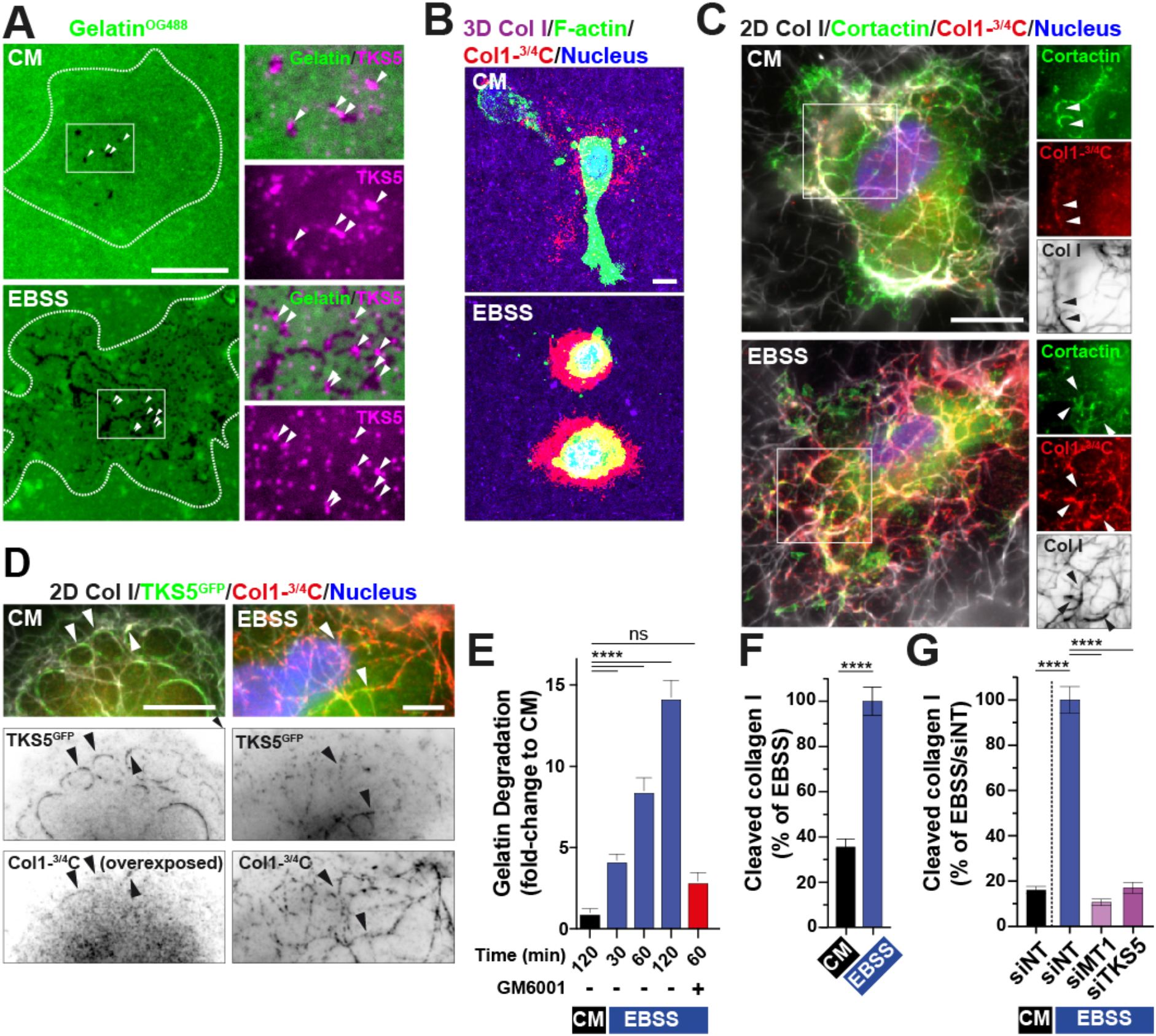
Enhanced collagenolysis and requirement for MT1-MMP and TKS5 in AA- and serum-starved cells. **(A)** MDA-MB-231 cells plated on fluorescently-labeled gelatin (green) for 60 min in CM or EBSS starvation medium depleted for AAs and serum and stained for TKS5 (magenta). Higher magnification of boxed regions is shown in the insets with an inverted lookup table for TKS5. Arrowheads, TKS5-positive invadopodia; dotted line, cell contour. Scale bar, 10 μm. **(B)** MDA-MB-231 cells embedded in a 3D collagen I gel (magenta) for 6 hrs in the indicated medium and stained for cleaved collagen (red); F-actin (green); nucleus (blue). Scale bar, 10 μm. **(C)** MDA-MB-231 cells cultured on a fibrillar type I collagen network (gray scale) for 60 min in indicated medium and stained for cortactin (green); cleaved collagen fibers (red); nucleus (blue). Scale bar, 10 μm. Higher magnification of boxed regions is shown in the insets with an inverted lookup table for collagen fibers. Arrowheads, cortactin-positive invadopodia. **(D)** MDA-MB-231 cells expressing TKS5^GFP^ cultured on a collagen network as in panel C. GFP (green); cleaved collagen fibers (red); nucleus (blue). Right rows, inverted lookup grayscale table showing TKS5^GFP^ and cleaved collagen. Arrowheads point to collagenolytic TKS5-positive invadopodia. Scale bar, 10 μm. **(E, F, G)** Proteolysis of the pericellular matrix by MDA-MB-231 cells cultured in the indicated conditions.

The association of the invadopodia components, cortactin and TKS5, with the cleaved fibers was visible irrespective of nutrient availability (Figure 1CD). As reported, silencing of MT1-MMP or TKS5 inhibited collagen cleavage in MDA-MB-231 cells in replete conditions (Supplemental Figure 1EF) ^16^. The contribution of MT1-MMP and TKS5 to starvation-induced collagenolysis was assessed in MDA-MB-231 and Bx-PC3 cells, which expressed similar levels of the main invadopodia components (Supplemental Figure 1D). Strikingly, depletion of MT1-MMP or TKS5 abolished collagenolysis in EBSS in both tumor cell lines (Figure 1G and Supplemental Figure 1G). Similarly, MT1-MMP was required for induction of gelatinolysis in EBSS (Supplemental Figure 1H). Collectively, these findings indicate that increased matrix degradation by AA-depleted cells is mediated by a MT1-MMP and TKS5-dependent mechanism in breast and pancreatic tumor cells.

### Induction of the MT1-MMP collagenolytic response upon starvation of TNBC PDX explants

To generalize these findings to a model closer to the human disease, we used cells obtained from breast cancer patient-derived xenografts (PDX) ^18,19^. Cells isolated from several independent PDXs were cultured *ex vivo* in a 3D type I collagen gel in AA-replete or depleted conditions and their collagenolytic activity was assessed (Figure 2A). PDX-derived cells were primarily composed of cytokeratin-8-positive carcinoma cells and fibroblasts, the latter being identified based on their characteristic spindle-shaped morphology (not shown). In five out of eight PDXs, overnight incubation in EBSS resulted in a strong induction of pericellular collagen degradation as compared to complete medium (Figure 2BC). Moreover, treatment with GM6001 inhibited collagenolysis in this setting (Figure 2C). Starvation of the PDXs correlated with reduced phosphorylated 4E-BP1 (Ser65) level as a proxy for mTORC1 kinase activity as compared to conditions of nutrient sufficiency (Figure 2C). Additionally, we found a correlation between the intensity of the starvation-induced collagenolytic response with MT1-MMP, and to some extent TKS5, expression levels in PDX (for instance compare the response of TKS5^Low^ and MT1-MMP^Low^ HBCx-66, HBCx-92 and HBCx-172 and MT1-MMP^High^ HBCx-4B and HBCx-60 PDXs, Figure 2DE).

**Figure 2.**
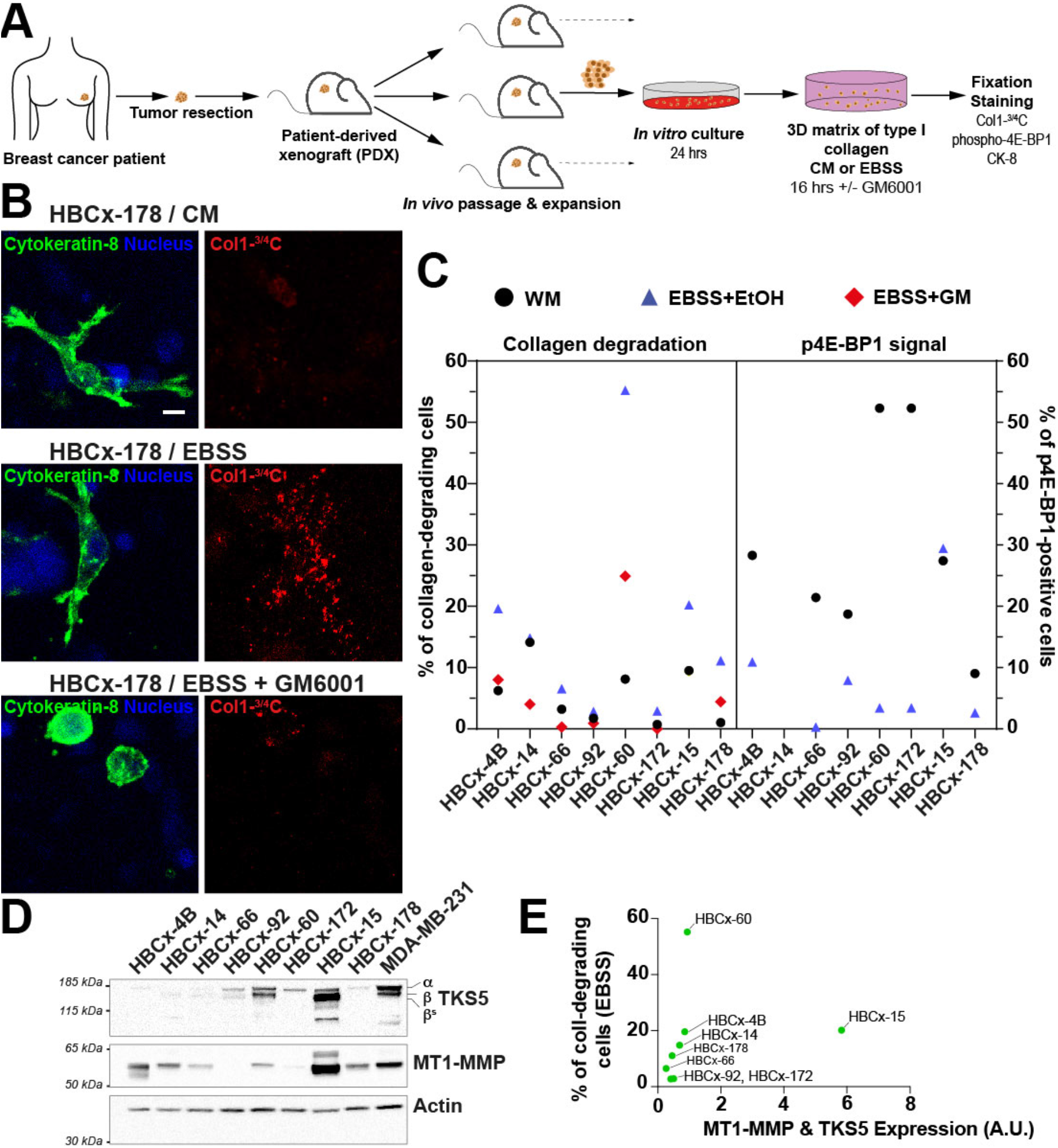
Induced collagenolysis in starved TNBC PDX ex vivo. **(A)** Scheme depicting the preparation and analysis of breast cancer derived PDX explants. **(B)** PDX cells embedded in type I collagen in the indicated culture conditions were fixed and epithelial breast tumor cells were stained for Cytokeratin-8 (green) and cleaved collagen (red). **(C**) Pericellular proteolysis of type I collagen (panel B) and levels of phosphorylated (p)4E-BP1 in and epithelial breast tumor cells (not shown) were plotted. Scale bar, 10 μm. **(D)** Representative western blots of TKS5 and MT1-MMP expression in the PDX-derived cells. Actin was used as a loading control. Molecular weights are in kDa. **(E)** Expression levels of MT1-MMP and TKS5 normalized to F-actin level based on the immunoblotting analysis show in panel D were summed up and plotted vs. the percentage of collagen degrading-cells for each PDX.

### Regulation of the collagenolytic response by mTOR signaling

mTOR is the master regulator of the cell’s response to nutrient and AA availability ^9^. As expected, mTORC1 activity was strongly repressed in cells cultured for 1 hr in EBSS as compared to CM as shown by the reduction in phosphorylated S6K (Thr389) and p4E-BP1 levels (Figure 3A and E). Replenishment of EBSS with free AAs similar to their concentration in CM (see Supplemental Table 4) partially restored mTORC1 activity (Figure 3A), and resulted in a 50-60% reduction of collagen cleavage by MDA-MB-231 cells as compared to EBSS (Figure 3B). These findings indicated that nutrient scarcity, in particular the lack of free AAs, strengthens the collagenolytic activity of breast MDA-MB-231 tumor cells, and that the absence of serum components accounted for 40-50% of the response. Along this line, it was recently shown that extracellular proteins including serum albumin can be internalized and catabolized in lysosomes and serve as an AA source to sustain cancer cells’ metabolic needs ^20–22^. We observed that supplementing EBSS with 3% bovine serum albumin (BSA) partially restored pS6K levels (Figure 3C, and supplemental Figure 2A) and resulted in a ~60% reduction of collagen cleavage as compared to EBSS alone (Figure 3D). All together, these data suggest a correlation between mTORC1 activity and ECM degradation, *i.e.* mTORC1 inhibition correlates with the induction of matrix degradation by breast tumor cells.

**Figure 3.**
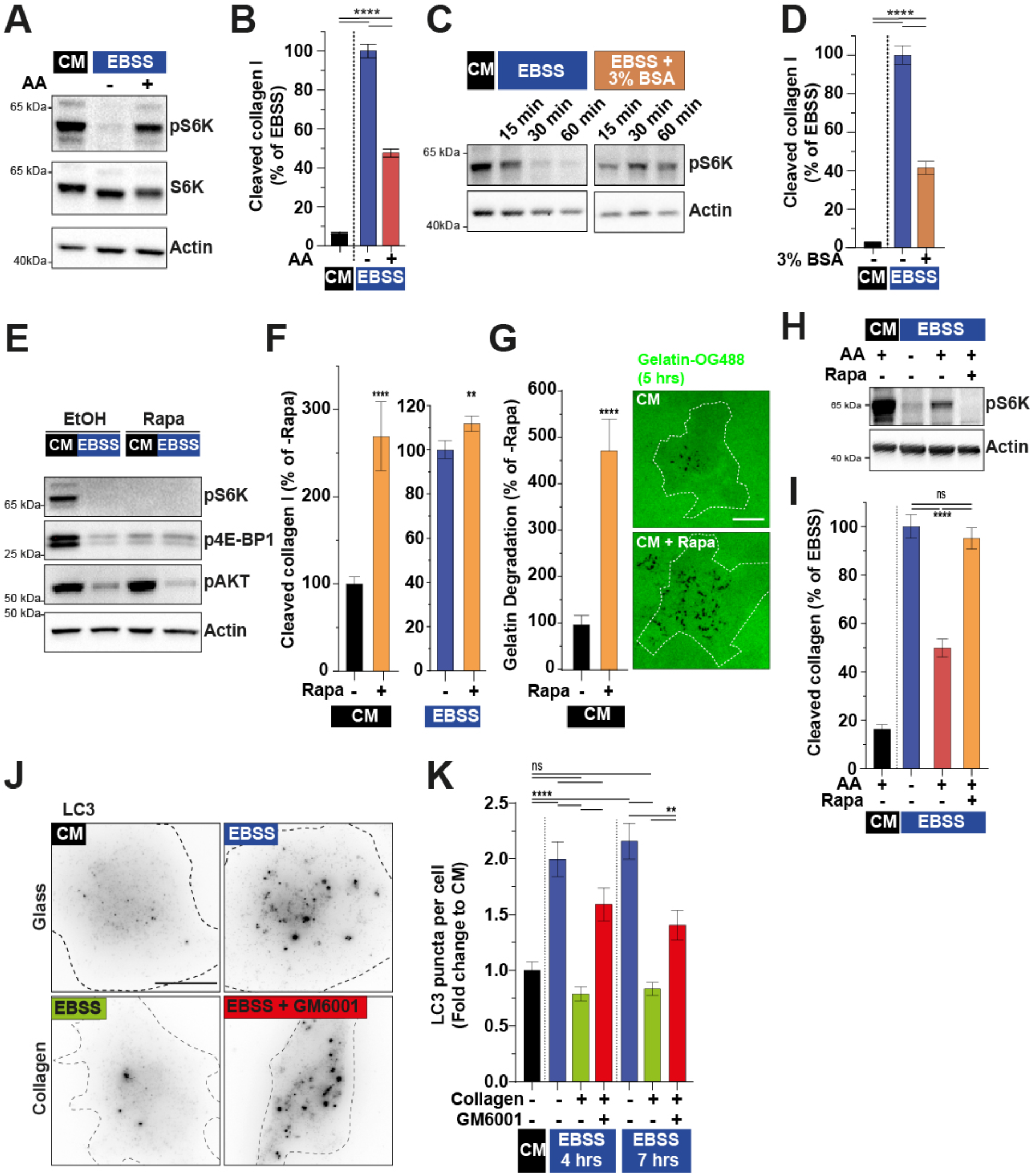
mTOR inhibition stimulates ECM degradation. **(A)** Total and phosphorylated S6K (Thr389) in MDA-MB-231 cells incubated for 60 min in indicated medium. Actin was used as a loading control. **(B)** Collagen cleavage by MDA-MB-231 cells incubated for 60 min in CM or EBSS supplemented with a complete AA mix. **(C)** Levels of pS6K in MDA-MB-231 cells incubated for 60 min in CM or EBSS medium supplemented with 3% BSA. **(D)** Collagen cleavage by MDA-MB-231 cells incubated for 60 min in the indicated medium. **(E)** Immunoblots of total and phosphorylated S6K, 4E-BP1 (Ser65) and AKT (Ser473) in MDA-MB-231 cells incubated for 60 min in the indicated medium in the presence of mTOR inhibitor, rapamycin or with corresponding vehicle with actin used as a loading control. **(F)** Collagen cleavage by MDA-MB-231 cells incubated for 60 min in indicated medium with or without rapamycin. **(G)** Gelatin degradation by MDA-MB-231 cells incubated for 5 hrs in CM with or without rapamycin. Scale bar, 10 μm. **(H, I)** Representative immunoblots of phosphorylated S6K (panel H). Collagen cleavage by MDA-MB-231 cells incubated for 60 min in CM or EBSS medium supplemented with AA in the presence or absence of rapamycin (panel I). **(J, K)** Quantification of autophagy LC3 puncta in MDA-MB-231 cells cultured in the indicated conditions normalized to the mean value in cells grown in CM medium on plastic ± SEM. Scale bar, 10 μm.

In agreement with this assumption, we observed that inhibition of mTORC1 activity by acute rapamycin treatment of cells grown in nutrient-replete conditions (CM) (Figure 3E, and Supplemental Figure 2B-D), resulted in a ~2.5-4-fold increase in collagen or gelatin degradation (Figure 3FG). In contrast, in EBSS, collagenolysis was only marginally increased upon rapamycin treatment (Figure 3F), in conjunction with the fully repressed mTORC1 status in starved cells (Figure 3E, and Supplemental Figure 2B-D). Interestingly, the inhibition of collagenolysis induced by supplementation of EBSS with free AAs was abolished in cells treated with rapamycin in parallel with mTORC1 repression (Figure 3HI). Thus, we conclude that the repression of mTORC1 activity in response to starvation is required for the induction of ECM degradation.

Autophagy is activated in response to starvation-induced mTORC1 repression ^23^. We analyzed the autophagy response of MDA-MB-231 cells and the influence of type I collagen exposure by staining for the autophagy marker, LC3. As expected, LC3-positive vesicular structures increased upon starvation of MDA-MB-231 cells cultured on plastic as compared to replete conditions. Autophagy was significantly reduced in starved cells cultured in the presence of type I collagen (Figure 3JK); in addition, this effect was partially abrogated by GM6001 (Figure 3JK). Collectively, our results suggest a novel feedforward loop mechanism in which AA scarcity represses mTORC1 activity, inducing ECM breakdown through a TKS5-, MT1-MMP-dependent mechanism that may produce AA resources ^6,7^, which, in turn, restore mTOR activity and downmodulate the autophagy response in tumor cells.

### Endocytic arrest and CCP retention of MT1-MMP in starved cells

Total levels of MT1-MMP remained steady for at least 6 hrs in starved cells suggesting some redistribution of a preexisting pool to support the increase in collagenolysis (Supplemental Figure 3A). The influence of nutrient availability on the distribution of MT1-MMP was analyzed. Confirming previous observations, MT1-MMP fused with a GFP variant (pHLuorin)-tag localized predominantly in perinuclear late endosomes/lysosomes ^24^ (Figure 4AB). In addition, in cells grown in replete conditions, MT1-MMP was detected in plasma membrane accumulations in association with the underlying collagen fibers (*i.e.* invadopodia, Figure 4A). In contrast, MT1-MMP redistributed to an extensive dotty-like surface pattern in cells cultured in EBSS (Figure 4B). Endocytic CCPs display an archetypical plasma membrane dotty distribution. Moreover, the LLY^573^ motif in the carboxy-terminal tail of MT1-MMP is known to interact with the clathrin adaptor AP-2 complex involved in MT1-MMP surface clearance (^25^ and see below). Counterstaining for the α-adaptin subunit of AP-2 revealed that MT1-MMP-positive puncta were in close proximity to CCPs in starved cells (Figure 4B). Additionally, we noticed a 1.6-fold increase in the density of CCPs at the plasma membrane of starved cells as compared to cells grown in CM (Figure 4C).

**Figure 4.**
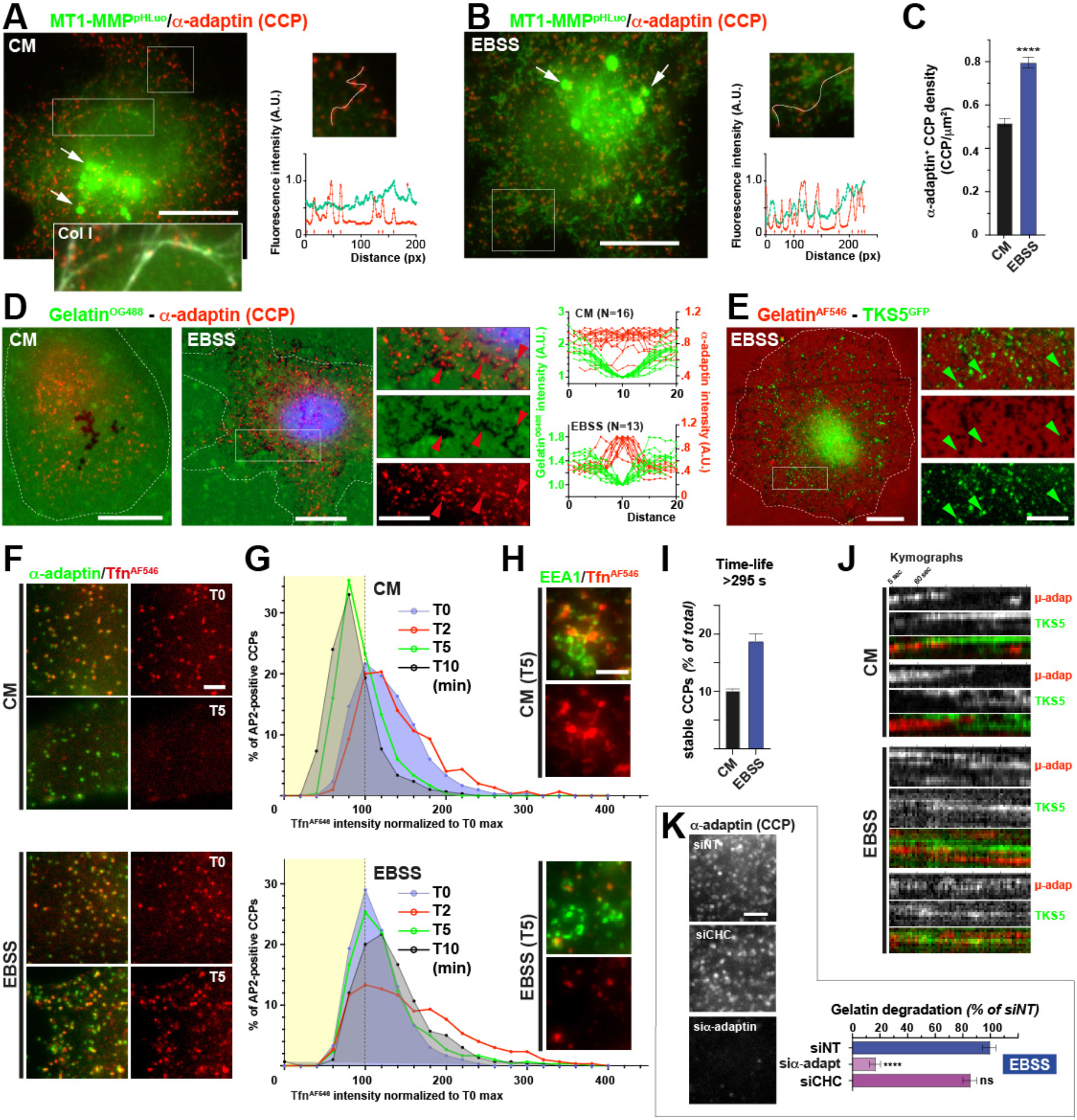
Starvation induces MT1-MMP endocytic arrest. **(A, B)** Distribution of MT1-MMP^pHLuorin^ (green) and α-adaptin-positive CCPs (red) in MDA-MB-231 cells cultured on fibrillar type I collagen in indicated medium. White arrows, fluorescence signal of MT1-MMP^pHLuorin^ in endolysosomes visible after cell fixation. Inset show higher magnification of the boxed region (type I collagen, gray). Scale bar, 10 μm. Right panels, intensity profile (linescan) of α-adaptin and MT1-MMP^pHLuorin^ signals (red arrows, CCPs). **(C)** Mean density of α-adaptin-positive CCPs ± SEM (CCP/μm^2^). **(D)** Distribution of α-adaptin-positive CCPs (red) in cells plated on fluorescently-labeled gelatin (green) in indicated medium. Scale bars, 10 μm; 2 μm (insets). CCP associated with gelatin degradation spots (red arrowheads). Right panels, linescan intensity profiles of α-adaptin (red) and gelatin fluorescence in degraded areas. **(E)** TKS5^GFP^-positive puncta associated with gelatin (red) degradation spots (green arrowheads). Scale bars, 10 μm; 2 μm (insets). **(F)** Fluorescence signal of Tfn^AF546^ (red) associated with α-adaptin-positive CCPs (green) in cells cultured in the absence of labeled Tfn in the indicated medium. Scale bar, 2 μm. **(G)** Kinetics of Tfn^AF546^ uptake in MDA-MB-231 cells incubated in the indicated medium. **(H)** Internalization of Tfn ^AF546^ (red) in EEA1-enriched endosomes (green) is reduced upon cell starvation. Scale bar, 2 μm. **(I)** Percentage of stable CCPs ± SEM (lifetime > 295 s). **(J)** Kymographs showing CCP and TKS5 dynamics in cells expressing μ-adaptin^mCh^ and TKS5^GFP^ plated on unlabeled gelatin in the indicated medium. Cells were imaged by TIRF-M every 5 s for 5 min. **(K)** Gelatinolysis by starved cells silenced for α-adaptin or CHC. Right panels show α-adaptin-positive CCPs in the different cell populations. Scale bar, 2 μm.

Similarly, we observed a striking association of gelatin degradation spots and AP-2-positive CCPs in cells plated on fluorescently-labeled gelatin matrix grown in EBSS medium (Figure 4D, right panel). In order to exclude that this association was by chance due to the high density of CCPs and degradation spots, AP-2 positions were randomly scrambled 5000 times, and AP-2 association with gelatin degradation was calculated for each scrambled image (Supplemental Figure 3BC). The observed association values exceeded all those calculated for randomly scrambled images ruling out that AP-2 association with degradation spots was random (Table S5). Moreover, MT1-MMP and TKS5 punctate accumulations also coincided with the dotty degradation pattern that appeared as a strong emerging feature of starved cells (Figure 4E, and Supplemental Figure 3DE and F).

Constitutive endocytosis of the transferrin-receptor (TfnR) is mediated by clathrin and AP-2 and can be readily monitored using fluorophore-conjugated Tfn. We followed the decay of Tfn^AF546^ from α-adaptin-positive CCPs overtime in cells cultured in CM or EBSS as a quantification of clathrin-mediated endocytosis (CME). While CCP-associated Tfn^AF546^ rapidly decayed in cells incubated in CM medium in the absence of fluorescent ligand, the intensity of receptor-bound Tfn^AF546^ associated with CCPs remained almost constant over the 10-min pulse in EBSS (Figure 4FG). Internalized Tfn^AF546^ rapidly reached EEA1-positive early endosomes in cells incubated in nutrient replete conditions, while the amount of Tfn^AF546^ detected in early endosomes was much lower in starved cells consistent with the reduction in Tfn uptake (Fig. 4H). Additionally, we followed the dynamics of μ-adaptin^mCh^-positive CCPs by TIRF-M and found a ~2-fold increase in the percentage of stable CCPs (lifetime > 295 s) in EBSS *vs*. CM conditions (Figure 4I and Movie S1), in agreement with the observed increase in CCP density and reduced CME flux. Interestingly, TIRF-M also revealed some association between CCPs and TKS5^GFP^-positive puncta that formed in their vicinity (Figure 4J and Movie S1). Similar to stable CCPs in starved cells, adjacent TKS5^GFP^-positive puncta also appeared to be long-lived (Figure 4J, and Movie S1). Finally, we found that under conditions of endocytic arrest in starved cells, the induction of gelatinolysis was abrogated by α-adaptin knockdown (Figure 4K and Supplemental Figure 3G). This was in sharp contrast to the silencing of clathrin heavy chain that did not significantly impair matrix degradation nor AP-2 cluster formation (Figure 4K). All together, these observations highlight the requirement for the clustering of surface-exposed MT1-MMP to sustain the starvation-induced ECM degradation response through a mechanism, which, likely, involves the interaction of MT1-MMP with AP-2 in arrested CCPs (see Supplemental Figure 3H, and ^25^).

The capacity of cells to remodel collagen fibers was compared in nutrient replete or deplete environments by live cell imaging. Movies confirmed that TKS5^GFP^-positive invadopodia on the ventral surface of cells in replete conditions were highly dynamic, rapidly expanded in size and pushed bundles of collagen fibers aside (Supplemental Figure 4A and Movie S2, ^13^). In contrast, punctate and mostly static TKS5-positive structures showing little expansion over time were visible in AA-starved cells (Supplemental Figure 4B and Movie S3). These observations demonstrate that tumor cells switch from a potent matrix remodeling and invasive mode in replete conditions typical of disseminating cells, to an exclusive matrix degradation and possibly nutrient sourcing program in nutrient-scarce conditions.

## Discussion

We show that depletion of extracellular AAs and serum to replicate conditions of nutrient scarcity in a collagen-rich microenvironment elicits a robust cancer cell-autonomous collagenolytic response, exceeding by one-order-of-magnitude the ECM-degradative activity of invasive breast and pancreatic cell lines and breast PDXs. The pericellular ECM-degradation response to starvation is triggered by mTOR inactivation and we identified the key invadopodia components, TKS5 and MT1-MMP, as major players. In contrast to its association to dynamically forming invadopodia at ECM contact sites typical of invasive cells under nutrient-replete conditions ^12,13^, surface-exposed MT1-MMP accumulates at arrested CCPs in cells in a nutrient-scarce environment. Intriguingly, we observed some association between dynamic TKS5-positive assemblies and CCPs, which is enhanced upon starvation. Interestingly, several CME regulators including inositol 5-phosphatase, SHIP2, and its product, phosphatidylinositol 3,4-bisphosphate (PI(3,4)P2), the F-BAR domain proteins, CIP4 and FBP17, the Arp2/3 complex activator, N-WASP, and cortactin are known TKS5 interactors involved in invadopodia formation, suggesting that related mechanisms operate at CCPs and invadopodia ^16,26–28^. Interaction of TKS5 with stable CCPs in conjunction with MT1-MMP clustering based on binding to the AP-2 clathrin-adaptor complex are probably key to the repurposing of CCPs into powerful ECM-degradative assemblies.

A dual role for collagenolytic invadopodia has been found during tumor cell invasion ^13^. On the one hand, limited proteolysis of individual collagen molecules by invadopodial MT1-MMP can soften the fibrils to facilitate cell passage during confined invasion ^13^. On the other hand, invadopodia can generate outward forces to push collagen fibers aside using the energy of actin polymerization ^13,29^. Although CCPs have been found to form in association with and can grab collagen fibers ^30^, actin-based forces generated at CCPs are inwardly oriented to facilitate budding of endocytic clathrin-coated vesicles ^31^. Thus, it is unlikely that CCPs could exert pushing forces on matrix fibers. It is more plausible that the ~10-fold increase in collagenolysis under nutrient restriction conditions leads to the fragmentation of ECM fibers facilitating their internalization and incorporation in the cell metabolism ^6–8^. It is thus tempting to speculate that MT1-MMP-rich CCPs in starved cells may have limited impact on invasion and cell movement, rather promoting a vigorous ECM degradation nutrient sourcing program.

Earlier studies in cell lines and in drosophila and mouse models revealed that genetic or pharmacological inhibition of mTOR kinase impedes endocytosis, similar to our observations ^32–35^. Other reports highlighted that nutrient scarcity and mTORC1 inhibition stimulate the nutritional use of extracellular proteins and that mTORC1/2 inhibition induces macropinocytosis, the main route for the uptake of extracellular proteins such as BSA by cancer cells ^7,20–22,36,37^. Collectively, despite mechanistic details that are missing, these data point to some opposite effects of starvation and mTOR inhibition on the downmodulation of CME and activation of macropinocytic (or related phagocytic) uptake, which could cooperate in the production and internalization of ECM fragments by tumor cells. Our study provides a novel example of the extreme capacity of cancer cells to rewire their nutritional plans and metabolism for survival and growth in adverse conditions by repurposing an ECM proteolysis machinery. It also underscores potential limitations of anti-mTOR therapeutic strategies as mTOR inhibition can unleash the ECM-degradative potential of carcinoma cells.

## Methods

### Cell culture, transfection and siRNA treatment

Human MDA-MB-231 breast adenocarcinoma cells obtained from ATCC (ATCC HTB-26) were grown in L-15 medium (Sigma-Aldrich) supplemented with 15% fetal calf serum (FCS) and 2 mM Gln (ThermoFisher Scientific) at 37°C in 1% CO_2_. The human pancreas adenocarcinoma cell line Bx-PC3 obtained from ATCC (ATCC CRL-1687) was grown in RPMI-1640 medium (ThermoFisher Scientific) supplemented with 10% fetal calf serum at 37°C in 5% CO_2_. Both cell lines were routinely tested for mycoplasma contamination. MDA-MB-231 cells stably expressing TKS5^GFP^ or MT1-MMP^pHLuorin^ were generated by lentiviral transduction ^13^. For transient expression, MDA-MB-231 cells were transfected with the plasmid constructs using AMAXA nucleofection (Lonza) and analyzed by live cell imaging 48 h after transfection. For starvation experiments, cells were cultured in EBSS medium (ThermoFisher Scientific) supplemented with MEM Vitamins (Gibco, composition in Supplemental Table 1) at 37°C in 5% CO_2_. For knockdown experiments, cells were treated with the indicated siRNAs (50 nM final concentration) using Lullaby (OZ Biosciences) according to manufacturer instructions and analyzed after 72 hrs of treatment. For silencing of α-adaptin and clathrin heavy chain (CHC), cells were treated twice at 48 hrs interval and analyzed 120 hrs after initial treatment. The siRNAs used for this study are listed in Supplemental Table 2.

### Antibodies and drugs

The source of commercial antibodies used for this study are listed in Supplemental Table 3. The source and working concentration of drugs used in this study are listed in Supplemental Table 4.

### Polymerization of type I collagen gel

A type I collagen polymerization mix was prepared on ice by adding 25 μM HEPES (final concentration) to a 2.2 mg/ml acidic-extracted type I collagen solution (Corning) and pH was adjusted to 7.5 with 0.34 N NaOH. When required for microscopic visualization of the collagen network, 2 to 5% of a ~2 mg/ml solution of AlexaFluor 647-conjugated type I collagen was added to unlabeled collagen in the polymerization mix. When required, drugs were added to the appropriate final concentration in the polymerization mix (see Supplemental Table 4). Polymerization was started by incubation at 37°C in a humidified chamber (CO_2_ cell incubator).

### Co-immunoprecipitation of MT1-MMP^phLuorin^-bound proteins

Cells stably expressing MT1-MMP^pHLuorin^ were plated on 100-mm dishes (3×10^6^ cells per dish). After 24 hrs cells were collected and lysed in lysis buffer (50 mM Tris-HCl, pH 7.5, 150 mM NaCl, 0.5 mM EDTA, 10 mM MgCl_2_, 10% glycerol, 60 mM β-glucoside, 1% NP-40, Phosphatase Inhibitor Cocktail 2 solution (1/100, SIGMA-Aldrich) and one tablet of cOmplete EDTA-Free protease inhibitor cocktail (SIGMA-Aldrich) for 15 min at 4°C under constant agitation. The lysate was centrifuged at 15,000 g for 10 min at 4°C, and the supernatant was incubated with 30 μl of control magnetic agarose beads (ChromoTek) for 30 min at 4°C under constant agitation. The precleared lysate was incubated with 30 μl of magnetic agarose beads coupled to anti-GFP nanobodies (GFPTrap; ChromoTek) for 1 hr at 4°C. GFP-Trap beads were washed with washing buffer A (lysis buffer without β-glucoside), followed by two washes with washing buffer B (without β-glucoside and NP-40). Beads were immediately heated at 95°C for 10 min in Laemmli buffer, stored at −20°C until Western blot analysis.

### Western blot analysis

RIPA sample buffer was prepared with 50 mM Tris-HCl pH8.0, 137 mM NaCl, 1% Triton X-100, 10 mM MgCl_2_, 10% glycerol and one tablet of cOmplete EDTA-Free protease inhibitor cocktail (SIGMA-Aldrich). For the study of phosphorylated proteins, RIPA buffer was supplemented with Phosphatase Inhibitor Cocktail 2 solution (diluted 1/100, SIGMA-Aldrich). A 24-well plate was layered with 20 μl of the 2.2 mg/ml unlabeled type I collagen polymerization mix as described above. After 3 min of polymerization at 37°C, the collagen gel was gently washed in PBS and a suspension of 1.5×10^4^ cells in CM or EBSS medium was added for 60 min at 37°C in 1% CO_2_ (CM) or 5% CO_2_ (EBSS medium) cell incubator. Cells were lysed in RIPA buffer for 30 min at 4°C and lysate was centrifuged for 30 min at 4°C at 13,000 rpm to remove debris and insoluble material. Proteins were separated by SDS-PAGE and transferred on a nitrocellulose membrane using the iBlot2 Dry Blotting System (Invitrogen). After incubating the membranes in 5% BSA or 5% skimmed milk in TBS-Tween 1%, proteins were detected by immunoblotting analysis with the indicated antibodies. Antibodies were visualized using the ECL detection system (GE Healthcare).

### Quantification of pericellular collagenolysis

To measure pericellular collagenolysis on a thin layer of type I collagen gel, a 18-mm diameter glass coverslip was layered with 200 μl of the ice-cold 2.2 mg/ml AlexaFluor 647(AF^647^)-labeled type I collagen polymerization mix as described above. Excess collagen solution was removed by pipette aspiration to leave a thin smear of collagen solution on the glass coverslip. After 3 min of polymerization at 37°C, the collagen gel was gently washed in PBS and 7×10^4^ cells were added and incubated for 1 to 7 hrs at 37°C in CM or EBSS medium in the presence or in the absence of AA supplements or drugs as indicated. Cells were pre-extracted with 0.1% Triton X-100 in 4% PFA in PBS for 90 sec at 37°C and fixed in 4% PFA in PBS for 20 min at 37°C. Coverslips were treated with 1% BSA in PBS for 30 min at room temperature then incubated with Col1-¾C and anti-cortactin antibodies diluted in 1% BSA in PBS for 2 hrs at 4°C. After three washes with PBS at 4°C, samples were counterstained with Cy3-conjugated anti-rabbit IgG and A488-conjugated anti-mouse IgG antibodies for 60 min at 4°C, extensively washed in PBS and mounted in Prolong-DAPI mounting medium (Invitrogen). Images were acquired with a wide-field microscope (Eclipse 90i Upright; Nikon) using a 100x Plan Apo VC 1.4 oil objective and a cooled interlined charge-coupled device (CCD) camera (CoolSnap HQ^2^; Roper Scientific). A z-dimension series of images was taken every 0.2 μm by means of a piezoelectric motor (Physik Instrumente). The system was steered by Metamorph software.

For quantification of pericellular collagenolysis in a 3D collagen network, 40 μl of a 6×10^4^ cells/ml cell suspension in the 2.2 mg/ml type I collagen polymerization mix was added on top of a 12-mm diameter glass coverslip and polymerization was performed for 30 minutes at 37°C. The indicated culture medium was added and samples were incubated for 6 hrs at 37°C. Samples were fixed, permeabilized and stained with Col1-¾C antibody as described above except that samples were counterstained with Phalloidin-Alexa488 to visualize cell shape. Image acquisition was performed with an A1R Nikon confocal microscope with a 40x NA 1.3 oil objective using high 455 sensitivity GaASP PMT detector and a 595 +/− 50 nm band-pass filter. Quantification of Col1-¾C signal (cleaved collagen) was performed with a homemade ImageJ macro. Acquired z-planes were projected using maximal intensity projection in Fiji and Col1-¾C signal was determined using the thresholding command excluding regions <50-px to avoid non-specific signal. Col1-¾C signal area was normalized to the total cell surface (thin layer) or to the number of nuclei in field (3D network) and values normalized to control cells.

### Fluorescent gelatin degradation assay

MDA-MB-231 cells were plated for 1 to 5 hrs on Oregon Green 488 (OR^488^) or AF^594^-conjugated cross-linked gelatin (Invitrogen) in EBSS or CM medium in the presence or absence of rapamycin as previously described ^38^. Cells were pre-extracted with 0.1% Triton X-100 in 4% PFA in PBS for 90 sec at 37°C and fixed in 4% PFA in PBS for 20 min at 37°C and then stained with the indicated antibodies or with fluorescently-labeled phalloidin to stain F-actin. Cells were imaged with a 100x objective on a wide-field microscope equipped with a piezoelectric motor as above. For quantification of degradation, the area of degraded matrix (black pixels) measured with the threshold command of ImageJ was divided by the total cell surface and values were normalized to control cells. The regions of interest delimiting the gelatin degradation were saved for further analysis, such as the assessment of AP-2 association (see below). Linescans were performed using Fiji software.

### Quantification of CCP density

7×10^4^ MDA-MB-231 cells were plated on top of gelatin^OR488^ and incubated for 1 hr in CM or EBSS medium. After fixation and permeabilization, cells were stained with α-adaptin as described above. CCPs in the entire cell were detected using the Find Maxima command of ImageJ and the number of detected CCPs was divided by the area of the cell. CCPs positions were saved for further analysis (see below).

### Randomization of AP-2 distribution over gelatin degradation spots

To measure the association of α-adaptin positive CCPs with gelatin degradation spots, CCPs and degradation spots were detected as described above and their positions as well as the position of all pixels inside the cell (total pixels) defined by their X and Y coordinates were saved. For each CCP, (X, Y) positions were randomly drawn from all pixels of the cell, effectively changing the position of CCPs inside the cell in a random fashion (see supplemental Figure 3B). This randomization procedure was performed 5,000 times per cell and the number of CCPs associated with gelatin degradation was measured each time. The true value of CCP association with gelatin degradation was calculated and compared to the randomized values. Synthetic images displaying cell contour (white line), degradation spots (black) and associated CCPs (red crosses) were generated with ImageJ. This procedure was repeated for ten independent cells with *p*-values ranging from 0 to 0.0142 (mean *p*-value = 0.001).

### Tfn uptake assay

7×10^4^ MDA-MB-231 cells plated on a 18-mm diameter glass coverslip were incubated overnight at 37°C in CM. Cells were washed twice with PBS before incubation in EBSS or CM medium for 1 hr at 37°C, then transferred on ice and washed twice with ice-cold EBSS or L15 medium supplemented with 1% BSA and 20 mM HEPES pH 7.5. Coverslips were incubated with 20 μg/ml of AF^546^-conjugated Tfn (ThermoFisher) in the same medium for 1 hr at 4°C. Cells were fixed with 4% PFA in PBS or incubated in pre-heated CM or EBSS for 2, 5 or 10 min at 37°C before fixation. After permeabilization with 0.1% Triton X-100 in PBS for 15 min, samples were incubated with anti-α-adaptin (overnight at 4°C) or with anti-EEA1 antibodies (1 hr at room temperature), and then counterstained with AF488-conjugated anti-mouse antibodies (1 hr at room temperature). Stacks of images were acquired with a wide-field microscope (Eclipse 90i Upright; Nikon) steered by Metamorph software as described above. For analysis, the plane corresponding to the plasma membrane was selected. CCPs positive for α-adaptin in a selected region were detected and segmented using the manual threshold command of ImageJ. The regions of interest (ROI) were saved and copied on the Tfn image. The mean intensity of Tfn inside each ROI was measured and a frequency histogram was generated with a normalization to T0.

### Quantification of LC3-positive puncta

7×10^4^ MDA-MB-231cells were plated on collagen-coated or on non-coated 18-mm diameter glass coverslips as previously described and incubated for 4 to 7 hrs at 37°C in CM or in EBSS medium. Cells were fixed with 4% PFA in PBS for 10 min and permeabilized with 0.05% saponin (Sigma-Aldrich) in PBS for 10 min. Samples were blocked in PBS with 0.05% saponin and 5% FCS for 30 min at room temperature and stained with anti-LC3 and anti-p4E-BP1 antibodies for 2 hrs at room temperature. After three washes, samples were counterstained with Cy3-conjugated anti-mouse IgG and Alexa488-conjugated anti-rabbit IgG antibodies and mounted in Prolong-DAPI medium. Image acquisition was performed by wide-field microscopy as previously described. Quantification of LC3-positive vesicles was performed by maximal orthogonal projection of the series of optical sections (the distance between two sections is 0.2 μm). Cells were manually delimited using the p4E-BP1 signal while LC3 signal was denoised and thresholded to detect LC3-positive autophagic vesicles. Detected spots were counted and saved for visual verification. No manual correction was done. The average number of LC3-positive puncta per cell was normalized to the value in CM-treated cells set to 1.

### Dynamics of TKS5- and μ-adaptin-positive structures by live cell total internal reflection fluorescence microscopy (TIRF-M)

MDA-MB-231 cells transfected with GFP-tagged TKS5 and mCherry-tagged μ-adaptin were plated in CM or EBSS on glass bottom dishes (Ibidi Corporation) layered with unlabeled cross-linked gelatin as previously described. Simultaneous dual color TIRF-M sequences were acquired with an inverted microscope (Eclipse-Ti-E, Nikon) equipped with a 100x PlanApo TIRF objective (1.47 NA), a TIRF arm, an image splitter (DV; Roper Scientific) installed in front of the EMCCD camera (Photometrics) and a temperature controller. GFP and mCherry were excited with 491- and 561-nm lasers, respectively (50 mW, Gataca Systems) and fluorescent emissions were selected with bandpass and longpass filters (Chroma Technology Corp). The system was driven by Metamorph. For quantification of CCP dynamics, CCP lifetime was measured using the TrackMate plugin of FIJI ^39^.

At least 300 CCPs from at least 6 cells per condition and per experiment were tracked from three independent experiments. Data are expressed as mean lifetime ± sem.

### *Ex-vivo* culture of TNBC patient-derived xenografts

Breast cancer patient derived xenografts have been generated as described ^19^. After surgical excision of the tumor xenograft, tumor cells were dissociated in DMEM/F12 medium supplemented with collagenase and hyaluronidase (SIGMA-Aldrich, 1X final) in 10 mM HEPES, 7.5% BSA Fraction V (Gibco), 5 μg/ml insulin (SIGMA-Aldrich) and 50 μg/ml gentamycin (GIBCO) for 1 hr at 37°C on a rotating wheel at 180 rpm as previously described ^18^. Samples were washed with DMEM/F12 medium and digested with 0.25% of trypsin (Gibco) for 2 min at 37°C. Trypsin was neutralized in HBSS medium (Invitrogen) supplemented in 10 mM HEPES and 2% FCS. Then, samples were treated with dispase (5UI/mL, StemCell Technologies) and DNAse I (1mg/mL in DMEM, Sigma for 2 min at room temperature and then incubated in neutralization buffer supplemented with NH_4_Cl (0.8%, StemCell Technologies) to remove red blood cells. After filtration through a 40 μm Cell Strainer (Corning), tumor cells were plated in a 25-cm^2^ cell-culture flask for 16 hrs at 37°C in DMEM/F12 medium supplemented with 10% FCS. For the pericellular collagenolysis assay, non-attached PDX tumor cells in the culture supernatant were resuspended in a 2,2 mg/ml collagen I solution as described above and incubated for 16 hrs in CM or EBSS medium with or without GM6001. After fixation with PFA 4% for 20 min and permeabilization with Triton 0.1% in PBS for 5 min, samples were stained with Col1-¾C and anti-Keratin-8 (K8) antibodies (2 hrs at 4°C) or anti-phospho-4E-BP1 and anti-Keratin-8 antibodies (1 hr at 4°C), counterstained with fluorescently labeled secondary antibodies and mounted. Image acquisition was performed with an A1R Nikon confocal microscope as described above.

### Statistics and reproducibility

All data are presented as mean ± S.E.M. from at least three independent experiments except indicated otherwise. GraphPad Prism software was used for statistical analysis. Data were tested for normal distribution using the D’Agostino-Pearson normality test and nonparametric tests were applied otherwise. One-way ANOVA, Kruskal-Wallis or Mann-Whitney tests were applied as indicated in the figure legends and are summarized in Supplemental Table 5. Statistical significance was defined as *, *P*<0.05; **, *P*<0.01; ***, *P*<0.001; ****, *P*<0.00001; ns, not significant.

## Acknowledgements

The authors thank Ms Gaëlle Martin for expert technical expertise and Drs Harald Stenmark, Camila Raiborg and Stephen J. Weiss and members of PC’s lab for helpful comments during the preparation of this manuscript. C.C. was supported by a grant from the Cancéropôle Région Ile-de-France (2016-2-APD-04-ICR-1) to P.C. This work was supported by a donation from Mr Trond Paulsen (InvaCell Project), by Fondation ARC pour la Recherche contre le Cancer (PGA1 RF20170205408) and Fondation Ruban Rose (Prix Avenir 2018) and by institutional support from Institut Curie and Centre National pour la Recherche Scientifique to PC.

The authors declare no competing financial interests.

## Author contributions

Cecilia Colombero, David Remy: conceptualization, investigation, methodology, validation, formal analysis, manuscript review and editing. Sandra Antoine, Pedro Monteiro, Nadia ElKhatib: methodology, investigation, validation, analysis. Margot Fournier, Ahmed Dahmani, Elodie Montaudon: methodology. Guillaume Montagnac: supervision of TIRF-M analysis. Elisabetta Marangoni: supervision of PDX experiments. Philippe Chavrier: conceived the study, conceptualization, validation, visualization, original drafting and editing of the manuscript, funding acquisition, supervision and project administration.

