## Supplemental Figures and Tables for "mTOR repression in response to amino acid starvation promotes ECM degradation through MT1-MMP endocytosis arrest"

**Supplemental Figure legends**

**Supplemental Figure 1. AA starvation induces matrix degradation in pancreatic Bx-PC3 tumor cells. (A)** Pancreatic ductal adenocarcinoma Bx-PC3 cells were cultured on a layer of fibrillar type I collagen (gray) for 60 min in indicated medium and stained for cortactin (green) and cleaved collagen I (Col1-¾C, red). Scale bar, 10 µm. Insets, higher magnification of boxed regions using inverted lookup tables (collagen fibers are in blue, cortactin or Col1-¾C signal is in red). Scale bar, 5 µm. **(B)** Collagen cleavage by Bx-PC3 cells was measured by Col1-¾C neoepitope staining and normalized to mean value of cells starved in EBSS ± SEM. **(C)** Collagen cleavage by MDA-MB-231 cells incubated for 60 min in CM or EBSS medium supplemented with 15, 75 or 2000 ng/ml recombinant human TIMP2 protein from two independent suppliers (rhTIMP2#1 and #2). **(D)** Comparison of the expression of key invadopodia components by immunoblotting analysis in MDA-MB-231 and Bx-PC3 cell lysates with actin as loading control. Molecular weights are in kDa. **(E)** Representative western blots of MT1-MMP and TKS5 expression with actin as loading control in MDA-MB-231 cells treated with indicated siRNAs. **(F)** Collagen cleavage by MDA-MB-231 cells knocked-down for MT1-MMP or TKS5 or treated with a non-targeting siRNA and cultured in CM. **(G)** Collagen cleavage by Bx-PC3 cells knocked-down for MT1-MMP or TKS5 or treated with a non-targeting siRNA and cultured in EBSS medium. **(H)** MDA-MB-231 cells knocked-down for MT1-MMP or treated with a non-targeting siRNA were plated on fluorescently-labeled gelatin in CM or EBSS medium for 2 hrs. Cell shape was visualized by phalloidin staining. The black dotted line underlines the cell contour. Scale bar, 10 µm. The graph shows the gelatin degradation normalized to the degradation of cells grown in EBSS medium ± SEM.

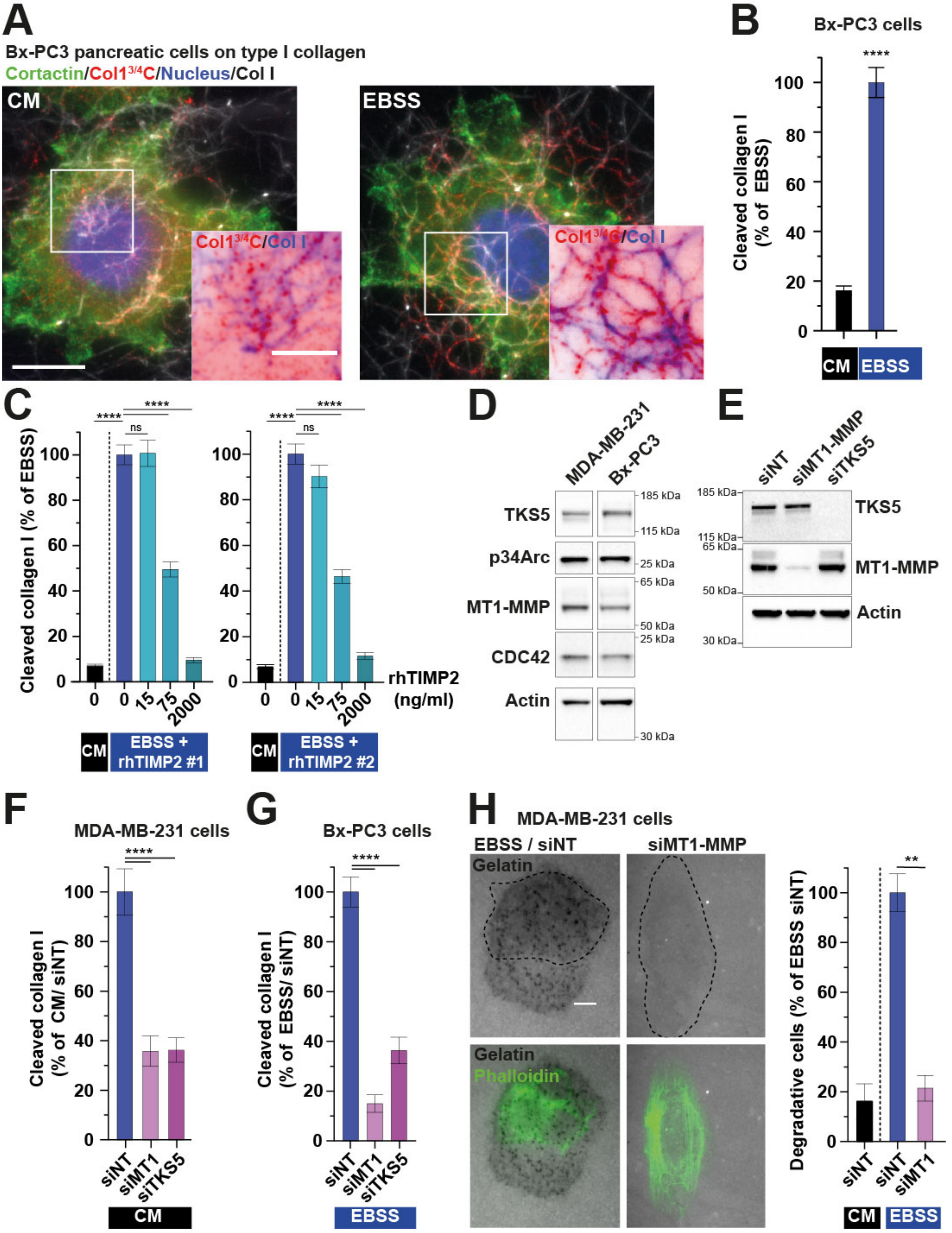
A
Bx-PC3 pancreatic cells on type I collagen
Cortactin/Col1 3/4C/Nucleus/Col I
CM
EBSS
Col1 3/4C/Col I
B
Bx-PC3 cells
Cleaved collagen I (% of EBSS)
****
CM
EBSS
C
Cleaved collagen I (% of EBSS)
ns
****
rhTIMP2 (ng/ml)
0
15
75
2000
CM
EBSS + rhTIMP2 #1
EBSS + rhTIMP2 #2
D
MDA-MB-231
Bx-PC3
TKS5
p34Arc
MT1-MMP
CDC42
Actin
185 kDa
115 kDa
25 kDa
65 kDa
50 kDa
30 kDa
E
siNT
siMT1-MMP
siTKS5
F
MDA-MB-231 cells
Cleaved collagen I (% of CM/ siNT)
siMT1
G
Bx-PC3 cells
Cleaved collagen I (% of EBSS/ siNT)
H
MDA-MB-231 cells
EBSS / siNT
siMT1-MMP
Gelatin
Phalloidin
Degradative cells (% of EBSS siNT)
**
siMT1

### SUP FIGURE 1

**Supplemental Figure 2. Phosphorylation of mTOR substrates in cells treated with rapamycin. (A)** Levels of phosphorylated S6K in MDA-MB-231 cells cultured in EBSS medium in the absence or presence of 3% BSA normalized to pSK6 levels in CM medium from two independent experiments (see Figure 3C). **(B-D)** Levels of phosphorylated S6K (panel A), 4E-BP1 (panel B) or AKT (panel C) normalized to actin levels in MDA-MB-231 cells cultured in CM or EBSS medium in the presence or absence of rapamycin from three independent experiments (see Figure 3E).

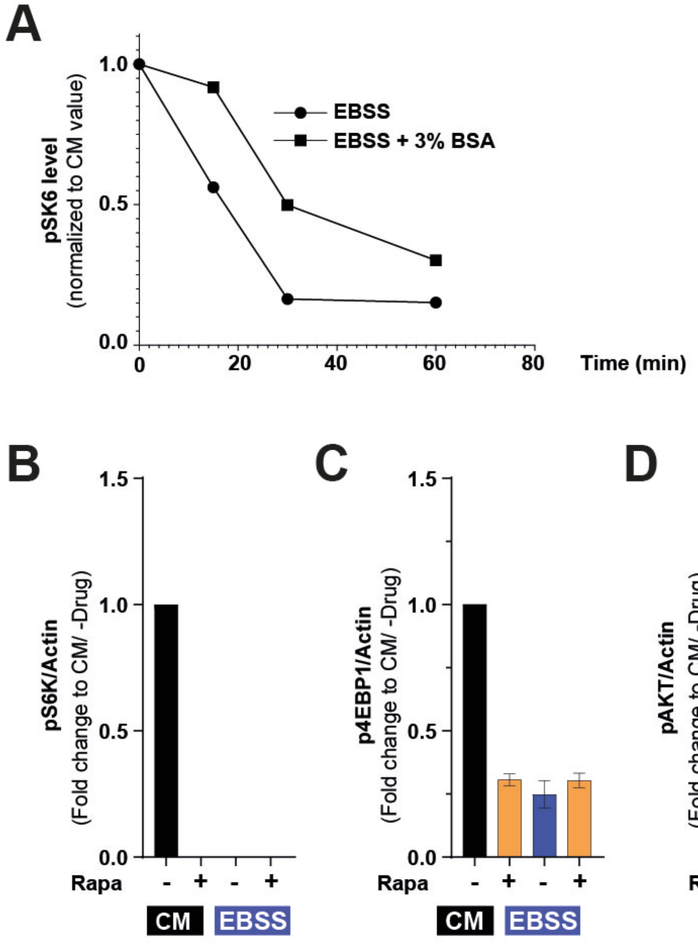


### SUPPLEMENTAL FIGURE 2

**Supplemental Figure 3. Matrix degradation correspond to MT1-MMP accumulation in arrested CCPs. (A)** MT1-MMP levels analyzed by western blot, normalized to actin in MDA-MB-231 cells incubated in the indicated medium for the indicated periods of time. Representative immunoblots are shown in the right panels. Molecular weight in kDa. **(B)** Synthetic images showing virtual CCPs (depicted as red crosses) overlapping with degraded areas (black). In the left image, the position of CCPs corresponds to their observed position in the original microscopy image (see Figure 4D, EBSS). The right image corresponds to one of the 5,000 scrambled images generated by randomization of CCP positions. **(C)** CCPs were scrambled 5,000 times and the histogram shows the number of randomized CCPs associated with gelatin degradation spots. The true unscrambled value (n=205) exceeds the randomized values, indicating high statistical confidence in non-random association of CCPs with degradation areas (see also Supplemental Table 5). **(D)** Synthetic images generated from the microscopy image in Figure 4E showing virtual TKS5$^{GFP}$-positive puncta (red crosses) superposed to degraded areas (black) as in panel B. (**E**) Histogram of the number of randomized TKS5 puncta associated with gelatin degradation spots as in panel C (see also Supplemental Table 5). **(F)** MDA-MB-231 cells expressing MT1-MMP$^{pHLuorin}$ (panel C) or TKS5$^{GFP}$ construct (panel D) were plated on AF$^{594}$-labeled gelatin for 60 min. White arrows, fluorescence signal of MT1-MMP$^{pHLuorin}$ in endolysosomes. Green arrowheads point to the accumulation of MT1-MMP$^{pHLuorin}$ in association with gelatin degradation areas. Scale bars, 10 µm. **(G)** Representative western blots of CHC, $\alpha$-adaptin or MT1-MMP expression with GAPDH as loading control in MDA-MB-231 cells treated with indicated siRNAs. Molecular weights are in kDa. Quantification of protein expression based on three ($\alpha$-adaptin and CHC) or two (MT1-MMP) independent experiments. (H) Lysates of MDA-MB-231 cells expressing

MT1-MMP$^{pHLuorin}$ were immunoprecipitated with GFP antibodies (GFP$^{Trap}$ IP). Total lysate before (input) and after immunoprecipitation (pIP) was loaded as control. Bound proteins were analyzed with MT1-MMP and α-adaptin antibodies Equal loading was controlled using GAPDH antibody.

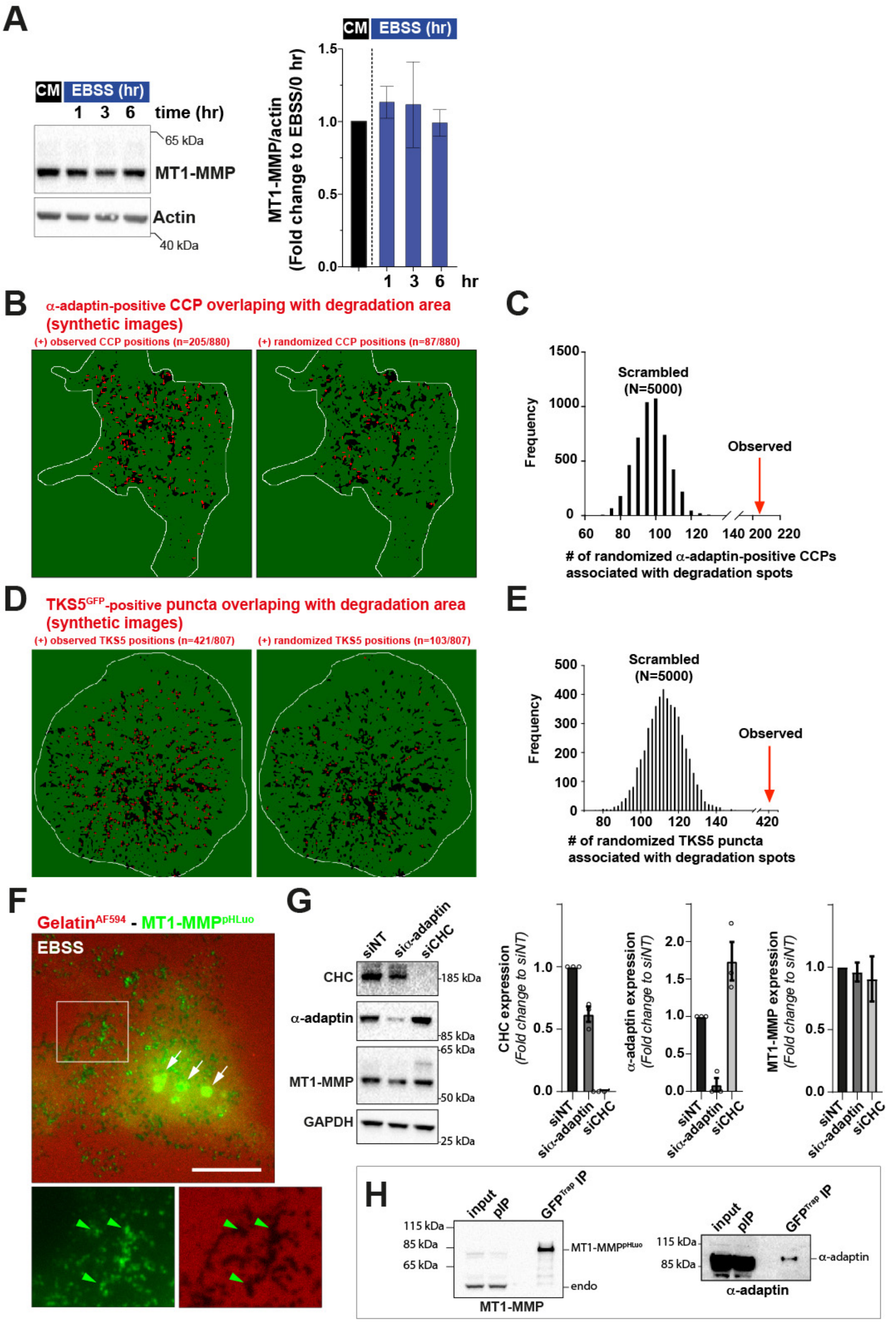

SUP FIGURE 3

**Supplemental Figure 4. Starvation induces invadopodia dynamics and matrix fiber remodeling program changes. (A, B)** MDA-MB-231 cells expressing TKS5$^{GFP}$ were cultured on fluorescently-labeled type I collagen in CM or EBSS and imaged over time (1 image/min) by video microscopy. The first and 20$^{th}$ images of representative time-lapse sequences are displayed with the upper row showing TKS5$^{GFP}$-positive invadopodia using an inverted grayscale lookup table, and the bottom row showing the collagen network using a Fire lookup table. The green dotted line underlines the cell contour. Color-coded time projections of seven images at 10-min intervals showing the dynamics of TKS5$^{GFP}$-positive invadopodia and the movement of type I collagen fibers over time.

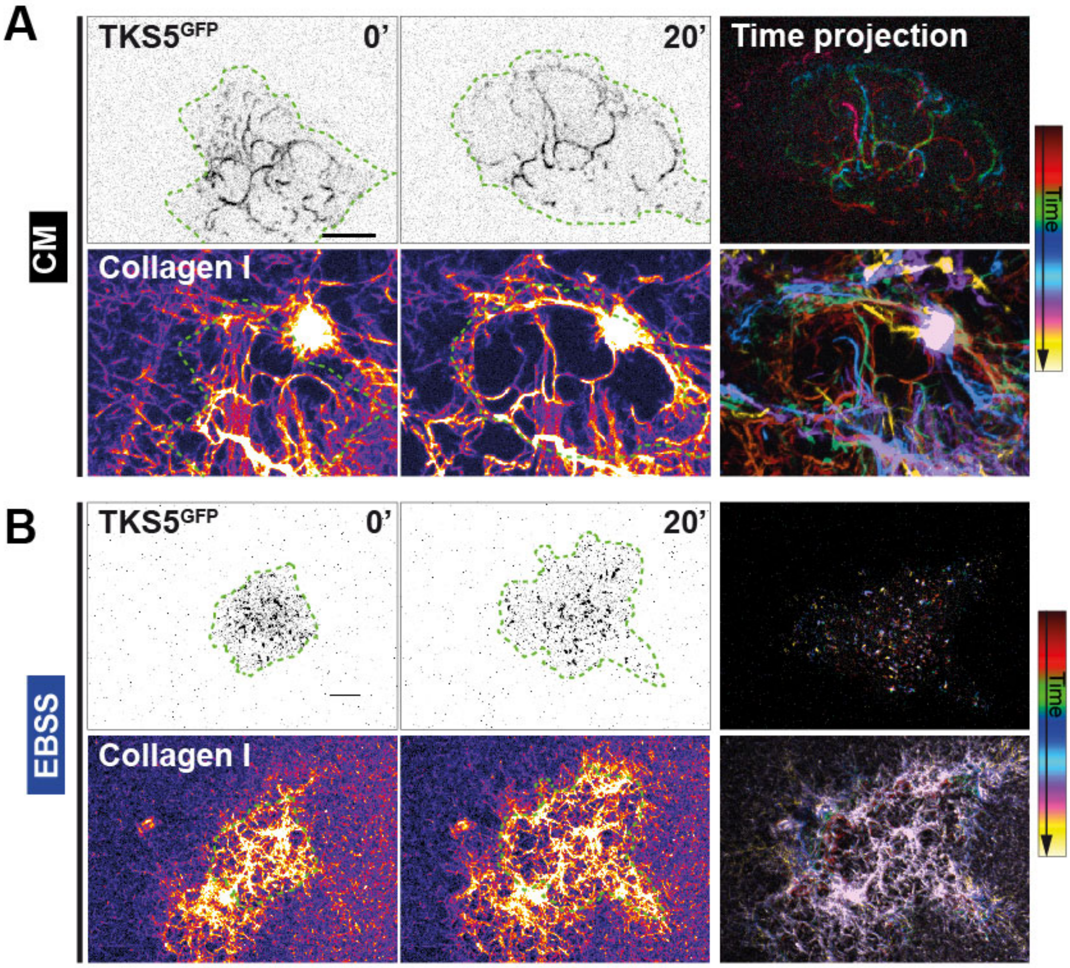


### SUP FIGURE 4

**Supplemental Movie legends**

**Supplemental Movie 1:** Shown is the dynamics of plasma membrane TKS5$^{GFP}$ (green) and CCPs labeled with μ-adaptin$^{mCherry}$ (red) in MDA-MB-231 cells plated on cross-linked gelatin in CM (upper panel) or EBSS medium (lower panel). Images were acquired every 5 s for 5 min by TIRF-M. In nutrient-replete conditions (CM), large and static TKS5$^{GFP}$-positive invadopodia form and TKS5 shows some transient association with CCPs, contrasting with long-lasting TKS5 interaction with CCPs observed in starved cells (yellow arrows). Scale bar, 2 µm.

**Supplemental Movie 2: TKS5 localizes to highly dynamic matrix fiber-remodeling elongated invadopodia in cells grown in nutrient-replete conditions.** Shown is a MDA-MB-231 cell expressing TKS5$^{GFP}$ (green). The cell is plated on a layer of fibrillar collagen (magenta) and is grown in nutrient-replete conditions. Images were acquired every 1 min for 60 min. The last sequence of the movie are still images of the time projection of seven frames separated by a 10-min interval with the different time points represented with the indicated pseudocolor coding. In nutrient-proficient medium, cells form dynamic elongated TKS5-positive invadopodia in association with the underlying collagen fibers that are actively remodeled.

**Supplemental Movie 3: TKS5 localizes to punctate, mostly static, structures in starved cells.** Shown is a MDA-MB-231 cell grown in nutrient-replete conditions and plated on a layer of fibrillar collagen (magenta). The cell expresses TKS5$^{GFP}$ (green). Images were acquired every 1 min for 60 min. The last sequence of the movie is a time projection of seven frames separated by a 10-min interval with the different time points represented with the indicated pseudocolor coding. Starved cells form static TKS5-positive puncta with minimal displacement of the underlying matrix fibers.

#### Supplemental Table 1. Medium composition

##### Vitamin mix

| Component | MEM Vitamin Solution 100 X (g/L) | L-15 Medium (g/L) |
|---|---|---|
| Choline Chloride | 0.1 | 0.001 |
| Folic Acid | 0.1 | 0.001 |
| Myo-Inositol | 0.2 | 0.002 |
| Niacinamide | 0.1 | 0.001 |
| D-Panthothenic Acid * ½Ca | 0.1 | 0.001 |
| Pirydoxal.HCl | 0.1 | 0.001 |
| Riboflavin | 0.01 | 0.0001 |
| Thiamine*HCl | 0.1 | 0.001 |

##### Amino acid mix composition

| Amino Acid | RPMI-1640 50 X (g/L) | L-15 Medium (g/L) |
|---|---|---|
| L-Alanine | - | 0.225 |
| L-Arginine (free base) | 10.0 | 0.5 |
| L-Asparagine | 2.84 | 0.25 |
| L-Aspartic Acid | 1.0 | - |
| L-Cystine | 2.5 | 0.12 |
| L-Glutamic Acid | 1.0 | 0.3 |
| Glycine | 0.5 | 0.2 |
| L-Histidine | 0.75 | 0.25 |
| Hydroxy-L-Proline | 1.0 | - |
| L-Isoleucine | 2.5 | 0.125 |
| L-Leucine | 2.5 | 0.125 |
| L-Lysine | 2.0 | 0.094 |
| L-Methionine | 0.75 | 0.075 |
| L-Phenylalanine | 0.75 | 0.125 |
| L-Proline | 1.0 | - |
| L-Serine | 1.5 | 0.2 |
| L-Threonine | 1.0 | 0.3 |
| L-Tryptophan | 0.25 | 0.02 |
| L-Tyrosine | 1.16 | 0.3 |
| L-Valine | 1.0 | 0.1 |

**Supplementary Table 2. siRNAs used in this study.**

| siRNA | Company | Targeted sequence (5'-...-3') |
|---|---|---|
| **MT1-MMP** *Smartpool* | Dharmacon L-004145-00-0005 | GGAUGGACACGGAGAAUUU<br>GGAAACAAGUACUACCGUU<br>GGUCUCAAAUGGCAACAUA<br>GAUCAAGGCCAAUGUUCGA |
| **TKS5** *Smartpool* | Dharmacon L-006657-00-0005 | ACAAUAACCUCAAAGAUGU<br>GGACGUAGCUGUGAAGAGA<br>CGACGGAACUCCUCCUUUA<br>GGAUAAGUUUCCCAUUGAA |
| **α-adaptin** | Merck Millipore | AAGAGCAUGUGCACGCUGGCCA |
| **Clathrin Heavy Chain (CHC)** | Elkhatib et al., Science 2017 | GCUGGGAAAACUCUUCAGATT |
| **Non-Targeting** | Dharmacon D-001810-01 | UGGUUUACAUGUCGUACUAA |

**Supplementary Table 3. Commercial antibodies and immunolabeling reagents used in this study.**

| Antigen | Company | Type (species) |
|---|---|---|
| **Collagen type I cleavage site (Col1–¾C)** *IF* | ImmunoGlobe (0217-050) | Polyclonal (Rabbit) |
| **Cortactin** *IF* | Merck (clone 4F11, 05-180) | Monoclonal (mouse) |
| **MT1-MMP (MMP14)** *IF, WB* | Merck (clone LEM-2/15.8, MAB3328) | Monoclonal (mouse) |
| **TKS5 (SH3PXD2A)** *IF, WB* | Novus Biologicals (NBP1-90454) | Polyclonal (rabbit) |
| **Paxillin** *IF* | BD Transduction Laboratories (610052) | Monoclonal (mouse) |
| **GFP** *IF* | Abcam (ab13970) | Polyclonal (chicken) |
| **LC3** *IF* | MBL (clone 4E12 M152-3) | Monoclonal (mouse) |
| **Alpha adaptin 2 (AP2)** *IF* | Abcam (ab2730) | Monoclonal (mouse) |
| **Alpha adaptin 2 (AP2)** *WB* | Abcam (ab2807) | Monoclonal (mouse) |
| **Anti Early Endosome Antigen 1 (EEA1)** | BD Transduction Laboratories (610457) | Monoclonal (mouse) |
| **Phospho-4E-BP1 (Ser65)** *IF, WB* | Cell Signaling (clone D9G1Q 13443S) | Polyclonal (rabbit) |
| **Phospho-(p70)S6 Kinase (Thr389)** *WB* | Cell Signaling (9205) | Polyclonal (rabbit) |
| **Phospho-AKT (Ser473)** *WB* | Cell Signaling (4060) | Polyclonal (rabbit) |
| **Total (p70)S6 Kinase** *WB* | Cell Signaling (clone 49D7 2708) | Monoclonal (rabbit) |
| **Total AKT** *WB* | Cell Signaling (9272) | Polyclonal (rabbit) |
| **β1 integrin** *WB* | Gift from C. Albiges-Rozo | Polyclonal (rabbit) |
| **Cytokeratin-8** *IF* | DSHB | Monoclonal (rat) |
| **Actin** *WB* | Sigma-Aldrich (clone AC-15 A1978) | Monoclonal (mouse) |
| **HRP-conjugated anti-rabbit IgG** | Sigma (A0545) | Goat |
| **HRP-conjugated anti-mouse IgG** | Jackson ImmunoResearch (115-035-062) | Goat |
| **Alexa Fluor 488 Phalloidin** | Molecular Probes (A12379) | |
| **Anti-rabbit Alexa488** | Molecular Probes (A21206) | Goat |
| **Anti-rabbit Cy3** | Molecular Probes (A21206) | Donkey |
| **Anti-chickenAlexaFluor488** | Molecular Probes (A11039) | Donkey |
| **Anti-mouse Cy3** | Jackson ImmunoResearch (715-165-151) | Donkey |
| **Anti-rat Alexa488** | Molecular Probes (A21208) | Donkey |

**Supplementary Table 4. Chemicals and reagents used in this study**

| Reagent | Company | Reference | Vehicle | Dilution |
|---|---|---|---|---|
| **RPMI 1640 amino acids solution** | Sigma-Aldrich | R7131 | Medium | 1/100 |
| **Bovine serum albumin solution 30%** | ThermoFischer Scientific | A7284 | Medium | 3% |
| **GM6001** | Merck Millipore | CC1100 | Ethanol | 40 µM |
| **Rapamycin** | Tocris Biotechne | 1292 | Ethanol | 20 nM |
| **Recombinant human tissue inhibitor of metalloproteinase (rhTIMP-2)** | - R&D Systems (rhTIMP-2#1)<br>- Sigma-Aldrich (rhTIMP-2#2) | 971-TM-010<br><br>SRP3174 | Medium | 15, 75 or 2000 ng/mL |
| **Transferrin from human serum, Alexa Fluor™ 546 Conjugate** | Invitrogen | 11530766 | Medium | 20 µg/ml |

**Supplementary Table 5. Analyzed variables and statistics used in this study.**

| Figure | Conditions | Mean | SEM | n | N | p value | Test |
|---|---|---|---|---|---|---|---|
| 1-E **Gelatin degradation** *(fold change to CM)* | **CM** | **1** | **0.2** | **42** | 2 | **-** | K-W |
| | EBSS 30 min | 4.2 | 0.4 | 42 | | <0.00001 | |
| | EBSS 60 min | 8.5 | 0.8 | 53 | | <0.00001 | |
| | EBSS 120 min | 14.2 | 1.1 | 42 | | <0.00001 | |
| | EBSS + GM6001 60 min | 2.9 | 0.6 | 42 | | ns | |
| 1-F **Cleaved collagen I** *(% of EBSS)* | CM | 35.7 | 3.5 | 59 | 3 | <0.00001 | M-W |
| | **EBSS** | **100** | **6.2** | **54** | | - | |
| 1-G **Cleaved collagen I** *(% of EBSS/siNT)* | CM siNT | 16.1 | 1.0 | 57 | 3 | <0.00001 | K-W |
| | **EBSS siNT** | **100** | **5.9** | **63** | | - | |
| | EBSS siMT1 | 10.6 | 1.4 | 62 | | <0.00001 | |
| | EBSS siTKS5 | 17.0 | 2.5 | 59 | | <0.00001 | |
| 3-B **Cleaved collagen I** *(% of EBSS/-AA)* | CM | 6.4 | 0.5 | 166 | 6 | <0.00001 | K-W |
| | **EBSS** | **100** | **3.5** | **171** | | - | |
| | EBSS+AA | 47.7 | 2.1 | 148 | | <0.00001 | |
| 3-D **Cleaved collagen I** *(% of EBSS/-BSA)* | CM | 2.6 | 0.4 | 91 | 3 | <0.00001 | K-W |
| | **EBSS** | **100** | **4.8** | **95** | | - | |
| | EBSS+BSA | 41.5 | 3.3 | 95 | | <0.00001 | |
| 3-F **Cleaved collagen I** *(% of - Rapa)* | **CM** | **100** | **8.0** | **90** | 4 | - | M-W |
| | CM + Rapa | 269.4 | 39.6 | 92 | | <0.00001 | |
| | **EBSS** | **100** | **4.0** | **64** | 3 | - | M-W |
| | EBSS + Rapa | 111.8 | 3.4 | 67 | | 0.0065 | |
| 3-G **Gelatin Degradation** (*fold change to CM)* | **CM** | **100.0** | **17.4** | **53** | 2 | - | M-W |
| | CM + Rapa | 472.0 | 68.0 | 54 | | <0.0001 | |
| 3-I **Cleaved collagen I** (*% of EBSS)* | CM | 16.3 | 1.1 | 87 | 4 | <0.0001 | K-W |
| | **EBSS** | **100** | **4.1** | **90** | | - | |
| | EBSS + AA | 53.8 | 3.1 | 93 | | <0.0001 | |
| | EBSS + AA + Rapa | 97.0 | 3.9 | 98 | | ns | |
| 3-K **LC3 puncta per cell** *(Fold change to CM)* | **CM - CollagenI** | **1.0** | **0.1** | **78** | 3 | - | K-W |
| | EBSS 4hrs - CollagenI | 2.0 | 0.2 | 102 | | <0.00001 | |
| | EBSS 4hrs +CollagenI | 0.8 | 0.1 | 87 | | ns | |

| | | | | | | | |
|---|---|---|---|---|---|---|---|
| | EBSS 4hrs +CollagenI +GM6001 | 1.6 | 0.1 | 88 | | NA | |
| | EBSS 7hrs - CollagenI | 2.1 | 0.2 | 104 | | <0.00001 | |
| | EBSS 7hrs +CollagenI | 0.8 | 0.1 | 101 | | ns | |
| | EBSS 7hrs +CollagenI +GM6001 | 1.4 | 0.1 | 94 | | NA | |
| 4-C **alpha-adaptin+ CCP density (CCP/µm²)** | **CM** | **0.52** | **0.02** | **41** | 4 | - | M-W |
| | EBSS | 0.8 | 0.02 | 60 | | <0.0001 | |
| 4-G **% of AP2-positive CCPs (CM)** | T0 | 100 | NA | 4459 | 3 | NA | NA |
| | T2 | 120 | | 2955 | | | |
| | T5 | 80 | | 1923 | | | |
| | T10 | 80 | | 1925 | | | |
| 4-G **% of AP2-positive CCPs (EBSS)** | T0 | 100 | NA | 5827 | 3 | NA | NA |
| | T2 | 100 | | 3614 | | | |
| | T5 | 100 | | 3776 | | | |
| | T10 | 120 | | 2121 | | | |
| 4-I **Stable CCPs** *(% of total)* | CM | 10.2 | 0.2 | 607 | 2 | NA | NA |
| | EBSS | 19.9 | 1.3 | 365 | | | |
| 4-K **Gelatin Degradation** *(% of EBSS siNT)* | **EBSS siNT** | **100** | **5.0** | **54** | 3 | - | K-W |
| | EBSS siα-adaptin | 17.0 | 4.1 | 50 | | <0.0001 | |
| | EBSS-siCHC | 85.6 | 5.1 | 48 | | n.s. | |

| Supplemental Figure | Condition | Mean | SEM | n | N | p value | Stat. |
|---|---|---|---|---|---|---|---|
| S1-B **Cleaved collagen I** *(% of EBSS)* | CM | 16.3 | 1.8 | 67 | 3 | <0.00001 | M-W |
| | **EBSS** | **100** | **6.1** | **88** | | - | |
| S1-C **Cleaved collagen I** *(% of EBSS/- rhTIMP2) Left panel* | CM | 6.9 | 0.9 | 46 | 3 | <0.00001 | K-W |
| | **EBSS** | **100** | **4.4** | **85** | | - | |
| | EBSS+15 ng/mL | 100.6 | 5.8 | 82 | | ns | |
| | EBSS+75 ng/mL | 49.5 | 3.3 | 80 | | <0.00001 | |
| | EBSS+2000 ng/mL | 9.5 | 1.2 | 77 | | <0.00001 | |
| S1-C **Cleaved collagen I** *(% of EBSS/ - rhTIMP2) Right panel* | CM | 6.9 | 0.9 | 46 | 2 | <0.00001 | K-W |
| | **EBSS** | **100** | **4.4** | **85** | | - | |
| | EBSS+15 ng/mL | 90.2 | 5.0 | 49 | | ns | |
| | EBSS+75 ng/mL | 46.4 | 3.0 | 52 | | <0.00001 | |
| | EBSS+2000 ng/mL | 11.6 | 1.5 | 41 | | <0.00001 | |
| S1-F **Cleaved collagen I** *(% of CM/ siNT)* | **CM siNT** | **100** | **9.4** | **57** | 3 | - | K-W |
| | CM siMT1 | 35.8 | 6.2 | 43 | | <0.00001 | |
| | CM siTKS5 | 36.2 | 4.9 | 45 | | <0.00001 | |
| S1-G **Cleaved collagen I** *(% of EBSS/ siNT)* | **EBSS siNT** | **100** | **6.0** | **96** | 3 | - | K-W |
| | EBSS siMT1 | 15.0 | 3.5 | 84 | | <0.00001 | |
| | EBSS siTKS5 | 36.3 | 5.3 | 70 | | <0.00001 | |
| S1-H **Degradative cells** *(% of EBSS/ siNT)* | CM siNT | 16.1 | 1.5 | 43 | 3 | NA | M-W |
| | **EBSS siNT** | **100** | **5.9** | **63** | | - | |
| | EBSS siMT1 | 10.6 | 1.4 | 62 | | 0.0015 | |
| S2-A **pS6K level** (*normalized to CM value)* | EBSS T0 | 1.0 | NA | NA | 2 | NA | NA |
| | EBSS T15 | 0.56 | | | | | |
| | EBSS T30 | 0.16 | | | | | |
| | EBSS T60 | 0.15 | | | | | |
| | EBSS + BSA 3% T0 | 1.0 | | | | | |
| | EBSS + BSA 3% T15 | 0.92 | | | | | |
| | EBSS + BSA 3% T30 | 0.50 | | | | | |
| | EBSS + BSA 3% T60 | 0.30 | | | | | |
| S2-B **pS6K/Actin** *(Fold change to CM/ -Drug)* | **CM** | **1** | **0** | NA | 3 | NA | NA |
| | Others | 0 | 4.0 | | | | |
| S2-C **p4E-BP1/Actin** (Fold change to CM/ -Drug) | **CM** | 1 | 0 | NA | 3 | NA | NA |
| | CM + Rapa | 0.3 | 0.02 | | | | |
| | EBSS | 0.2 | 0.05 | | | | |
| | EBSS + Rapa | 0.3 | 0.03 | | | | |
| S2-D **pAKT/Actin** *(Fold change to CM/ -Drug)* | **CM** | 1 | 0 | NA | 3 | NA | NA |
| | CM + Rapa | 1.3 | 0.4 | | | | |
| | EBSS | 0.2 | 0.06 | | | | |
| | EBSS + Rapa | 0.1 | 0.04 | | | | |
| S3-A **MT1-MMP/Actin** *(Fold change to EBSS/0 hr)* | **CM** | **1.0** | 0 | NA | 2 | - | K-W |
| | EBSS 1h | 1.0 | 0.1 | | | ns | |
| | EBSS 3hrs | 0.8 | 0.3 | | | ns | |
| | EBSS 6hrs | 0.9 | 0.1 | | | ns | |

| | Cell | Mode of randomized | Min-Max of randomized values | True value | p-value |
|---|---|---|---|---|---|
| S3-C **Randomization of AP2 positions** | 1 | 20 | 5-40 | 46/350 | 0 |
| | 2 | 41 | 16-63 | 91/641 | 0 |
| | 3 | 40 | 20-64 | 61/514 | 0 |
| | 4 | 50 | 27-74 | 127/442 | 0.0016 |
| | 5 | 40 | 20-65 | 54/394 | 0 |
| | 6 | 59 | 34-92 | 118/703 | 0.0142 |
| | 7 | 117 | 83-153 | 165/682 | 0 |
| | 8 | 98 | 63-133 | 204/880 | 0 |
| | 9 | 61 | 36-88 | 143/879 | 0 |
| | 10 | 51 | 30-77 | 112/575 | 0 |
| S3-E **Randomization of TKS5 positions** | 1 | 225 | 195-264 | 323/733 | 0 |
| | 2 | 174 | 140-214 | 308/804 | 0 |
| | 3 | 445 | 64-120 | 218/468 | 0 |
| | 4 | 555 | 495-630 | 670/435 | 0 |
| | 5 | 243 | 201-288 | 518/824 | 0 |
| | 6 | 158 | 124-200 | 349/1038 | 0 |
| | 7 | 94 | 68-124 | 335/886 | 0 |
| | 8 | 46 | 28-64 | 160/325 | 0 |
| | 9 | 112 | 80-142 | 421/807 | 0 |
| | 10 | 68 | 44-88 | 210/605 | 0 |

| Supplemental Figure | Condition | Mean | SEM | n | N | p value | Stat. |
|---|---|---|---|---|---|---|---|
| S3-G **CHC expression** (*Fold change to siNT*) | **siNT** | **1.0** | **0** | - | 3 | NA | NA |
| | siα-adaptin | 0.6 | 0.06 | | | | |
| | siCHC | 0 | 0 | | | | |
| S3-G **α-adaptin expression** (*Fold change to siNT*) | **siNT** | **1.0** | **0** | - | 3 | NA | NA |
| | siα-adaptin | 0.1 | 0.1 | | | | |
| | siCHC | 1.7 | 0.3 | | | | |
| S3-G **MT1-MMP expression** (*Fold change to siNT*) | **siNT** | **1.0** | **0** | - | 2 | NA | NA |
| | siα-adaptin | 0.9 | 0.07 | | | | |
| | siCHC | 0.9 | 0.2 | | | | |

S.E.M., standard error of the mean; n, sample number; N, number of independent experiments; ns, not significative; NA, not available.

Data were tested for normal distribution using the D'Agostino-Pearson normality test and nonparametric tests were applied otherwise.

Non-parametric tests: K-W, Kruskal-Wallis test; M-W, Mann-Whitney

Parametric tests: One-Way ANOVA

Statistical significance was defined as *, P <0.05; **, P <0.01; ***, P <0.001; ****, P <0.00001; ns, not significant.
